# A hypomorphic mutation in *Pold1* disrupts the coordination of embryo size expansion and morphogenesis during gastrulation

**DOI:** 10.1101/2022.03.07.483315

**Authors:** Tingxu Chen, Heather Alcorn, Sujan Devbhandari, Dirk Remus, Elizabeth Lacy, Danwei Huangfu, Kathryn V. Anderson

## Abstract

Formation of a properly sized and patterned embryo during gastrulation requires a well-coordinated interplay between cell proliferation, lineage specification and tissue morphogenesis. Following transient physical or pharmacological manipulations, pre-gastrulation stage mouse embryos show remarkable plasticity to recover and resume normal development. However, it remains unclear how mechanisms driving lineage specification and morphogenesis respond to defects in cell proliferation during and after gastrulation. Null mutations in DNA replication or cell-cycle related genes frequently lead to cell cycle arrest and reduced cell proliferation, resulting in developmental arrest before the onset of gastrulation; such early lethality precludes studies aiming to determine the impact of cell proliferation on lineage specification and morphogenesis during gastrulation. From an unbiased ENU mutagenesis screen, we discovered a mouse mutant, *tiny siren* (*tyrn*), that carries a hypomorphic mutation producing an aspartate to tyrosine (D939Y) substitution in Pold1, the catalytic subunit of DNA polymerase δ. Impaired cell proliferation in the *tyrn* mutant leaves anterior-posterior patterning unperturbed during gastrulation but results in an overall reduction in embryo size and in severe morphogenetic defects. Our analyses show that the successful execution of morphogenetic events during gastrulation requires that lineage specification and the ordered production of differentiated cell types occur in concordance with embryonic growth.

**Summary statement:** *Pold1* hypomorphic mutation caused reduced size and abnormal morphology of gastrulating mouse embryos, supporting the importance of coordinated embryo size, lineage specification and tissue morphogenesis for normal embryogenesis.

## Introduction

Gastrulation is a critical developmental process required for germ layer formation and the establishment of the body plan (Arnold and Robertson, 2009). Gastrulation initiates with the emergence of the primitive streak in the proximal posterior epiblast. As the streak extends to the distal tip of the embryo, epiblast cells undergo an epithelial mesenchymal transition (EMT) to form the mesoderm layer between the epiblast and the visceral endoderm (VE) (Kinder et al., 1999; Lawson, 1999; Lawson et al., 1991). Epiblast cells ingressing through the anterior region of the elongating primitive streak intercalate into the VE to form the definitive endoderm layer, which will give rise to the gut tube, and subsequently, the epithelium of endodermal organs, such as the pancreas and intestine (Kwon et al., 2008; Lawson et al., 1986; Lawson and Pedersen, 1987). Gastrulation requires tight spatiotemporal coordination of embryo size, tissue migration and cell fate determination. Despite extensive research on these topics, it has been challenging to untangle the complex interplay among these three key components. Previous studies used embryological methods in pre-implantation embryos to investigate size regulation during the pre- and early-gastrulation stages. These experiments found that double-sized embryos, formed by aggregating two 8-cell stage morula, underwent size regulation before gastrulation. The double-sized embryos showed an increase in cell-cycle length compared to controls; in addition, they lacked the proliferative burst that normally occurs before gastrulation. These two modes of regulating cell proliferation allowed the aggregated embryos to reach a normal size and cell number before E7.0 (Buehr and McLaren, 1974; Lewis and Rossant, 1982) and then to gastrulate. Conversely, undersized mouse embryos generated by removing one or two blastomeres from the 4-cell stage preimplantation embryo, sustained a prolonged proliferative burst, leading to an increase in cell number before the initiation of gastrulation (Power and Tam, 1993). Another study on size regulation in the mammalian embryo examined the response to reduced cell number in the early post-implantation embryo, an experimentally more refractory stage. Following treatment with mitomycin to inhibit cell proliferation, E7.0 embryos, with ∼80% of their cells eliminated, could still recover and complete gastrulation (Snow and Tam, 1979). These elegant studies suggest that intrinsic mechanisms operate within the pre- and early post-implantation embryo to monitor and control cell number both before and at the onset of gastrulation, supporting regulative development as an important feature of early mammalian embryogenesis. However, standard embryological and pharmacological methods are inadequate to further investigate how altered cell number in the gastrulating embryo impacts tissue patterning and morphogenesis. Therefore, to ask whether mechanisms of regulative development continue to act during gastrulation, it is crucial to explore alternative approaches to perturbing cell number.

Although many genes encoding cell cycle related proteins have been genetically inactivated to explore the effects of cell proliferation on embryo size and morphogenesis, the resulting phenotypes are generally not suitable for studies of gastrulation. For example, *Cyclin A2* (*Ccna2*) homozygous null mutants can be recovered only up to E5.5 (Murphy et al., 1997), whereas embryos lacking all D-type cyclins survive past gastrulation, with no overt phenotypes (Kozar et al., 2004). Another target for genetic perturbation of cell proliferation are the polymerases that replicate DNA. DNA Polymerase Delta (Pol δ), the subject of this report, plays multiple critical roles in DNA replication, with functions in DNA synthesis and repair (Jain et al., 2018). In mammalian cells, the Pol δ contains 4 subunits: the catalytic subunit, p125 (Pold1), and three regulatory subunits, p50 (Pold2), p66 (Pold3) and p12 (Pold4). Pold1 consists of two functional domains: a N-terminal 3’-5’ exonuclease with DNA proofreading activities and a C-terminal DNA polymerase that catalyzes DNA synthesis. Several mutant alleles have been generated for *Pold1*, but similar to targeted cell-cycle related genes mentioned above, the homozygous mutants either fail to survive beyond the onset of gastrulation or show no phenotypic defects during gastrulation. Null mutations in *Pold1* cause peri-implantation lethality (Uchimura et al., 2009). Two missense mutations have been reported for *Pold1*: a D400A substitution in the exonuclease domain and a L604K substitution in the DNA polymerase activity site (Fig. S1A) (Uchimura et al., 2009; Venkatesan et al., 2007). While developmentally normal, *Pold1*^*D400A/D400A*^ mice frequently died with swollen thymuses 3 months after birth (Uchimura et al., 2009). *Pold1*^*+/L604K*^ heterozygous mice underwent normal development but had a reduced lifespan and developed multiple tumor types, including lymphoma, adenoma and carcinomas of the liver and lung (Venkatesan et al., 2007). Although no specific phenotypes were reported for *Pold1*^*L604K/L604K*^ embryos, notably they died around E8.5, suggesting that missense mutations in the polymerase domain might perturb cell proliferation at a level compatible for investigation of gastrulation phenotypes.

In this study, we report a *Pold1* hypomorphic mutation identified in a phenotype-based genetic screen for recessive mutations causing gastrulation defects in mouse embryos. This mutation altered a conserved residue (D939Y) in the Pold1 DNA polymerase domain, caused reduced Pold1 protein expression, and resulted in compromised DNA synthesis. Mutant embryos could be retrieved up to E8.5; at this stage they were small with a siren-like morphology; hence we named the mutant *tiny siren* (*tyrn*). We investigated embryo growth and cell lineage differentiation in *tyrn* mutants at developmental stages between E6.5 and E8.5. The *tyrn* mutation impaired cell proliferation without affecting anterior-posterior patterning, but severely disrupted tissue morphogenesis during gastrulation. Our findings suggest normal cell proliferation is essential for mesoderm lineage allocation and is required to coordinate embryo size with cell movement for proper morphogenesis.

## Results

### *tyrn* mutant embryos show abnormal morphology but proper anterior-posterior patterning

To study the genetic regulation of gastrulation, we performed mouse ENU mutagenesis screens to uncover mutant embryos with abnormal morphology at E8.5 (Garcia-Garcia et al., 2005; Hernandez-Martinez et al., 2019; Huangfu et al., 2003; Migeotte et al., 2011). We isolated a mutant (later named *tyrn*) that exhibited not only a smaller overall size, but also a striking body shape and orientation (Fig. 1A) that was distinctly from other mutant phenotypes we had observed at this stage (Bazzi et al., 2017; Garcia-Garcia et al., 2005; Hernandez-Martinez et al., 2019; Huangfu et al., 2003; Migeotte et al., 2011; Zhou and Anderson, 2010). Instead of forming a U-shaped embryo with a well extended anterior-posterior (A-P) axis, the *tyrn* mutant embryos had a short A-P axis and lay relatively flat along the posterior side of the yolk sac, with the head misoriented towards the distal tip (Fig. 1A). To determine if the irregularly shaped mutants had established and correctly positioned the A-P axis, we performed whole mount *in situ* hybridizations (WISH) at E8.5 to a diagnostic set of markers. We found that although *tyrn* mutant embryos did not form a well-structured head, they did express *Foxg1* and *Otx2*, markers of forebrain and forebrain + midbrain, respectively, in discrete overlapping regions (Fig 1B-C). In addition, WISH detected relatively normal expression of *T* (*Brachyury*), which marks the posterior tail bud and notochord (Fig. 1D), and of *Foxa2*, which labels the floor plate of the neural tube, posterior notochord, notochordal plate and gut endoderm (Fig. 1E). Therefore, despite causing a highly unusual body shape, the *tyrn* mutation does not affect A-P patterning.

**Figure 1.**
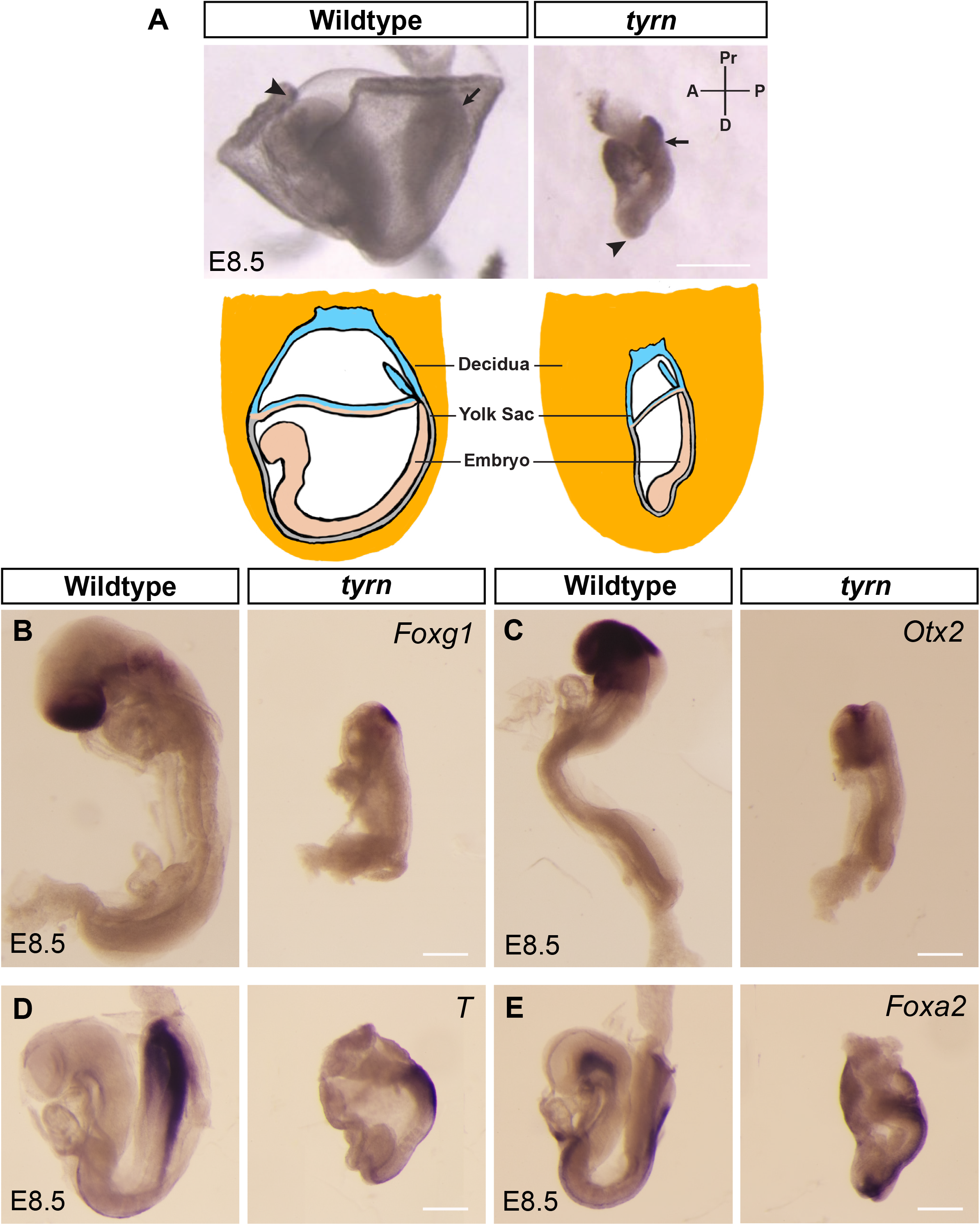
Characterization of *tiny siren* (*tyrn*), a mutant recovered from an ENU Screen. (A) Top: Wildtype and *tyr*n embryos recovered at E8.5. Embryos were aligned as they were in decidua. Mutant embryos showed a shifted body A-P axis, with a small, unstructured head located at the distal tip (arrowhead) and the tail at the posterior-proximal side (arrow). Scale Bar = 500 μm. Bottom: A cartoon that shows the orientation of wildtype and mutant embryos inside decidua. (B) *Foxg1* labels forebrain and (C) *Otx2* labels midbrain and forebrain in wildtype embryos. Mutant embryos expressed both *Foxg1* (B) and *Otx2* (C) in head. (D)*T* (*Brachyury)* marks the primitive streak and notochord in E8.5 wild types and *tyrn*. (E) Both wild types and *tyrn* expressed *Foxa2* in the floor plate, posterior notochord, notochordal plate and gut endoderm. n = 3 embryos per genotype. Scale Bar = 200 μm.

### *tyrn* is a hypomorphic allele of *Pold1*

To identify the causal mutation for the *tyrn* phenotype, we collected both wildtype and mutant embryos at E8.5 and performed whole-exome sequencing (Jain et al., 2017). We found a G to T transversion at nucleotide position 2815 of the *Pold1* open reading frame that generated an aspartate to tyrosine substitution (D939Y) within the DNA polymerase domain (Fig. 2A). The mutated aspartate residue is highly conserved across eukaryotic organisms from budding yeast to humans (Fig. 2B). Western blot analysis showed a reduced level of Pold1 in *tyrn* mutant embryos (Fig. 2C). To ask whether *Pold1* is the causative gene underlying the *tyrn* mutant phenotype, we performed a complementation test using the *Pold1*^*tm1b*^ null allele, derived from embryonic stem cells carrying a “knockout-first” tm1a allele (Skarnes et al., 2011) (Fig. 2D, Fig S1B). No *Pold1*^*tm1b/tm1b*^ embryos were recovered at post-implantation stages, consistent with the phenotype previously reported for *Pold1* null mutants (Uchimura et al., 2009). The *Pold1*^*tyrn/tm1b*^ embryos produced from a *Pold1*^*tm1b/+*^ and *tyrn/+* cross did not initiate gastrulation and failed to survive past ∼E7.5 (Fig. 2D), demonstrating that the *tm1b* and *tyrn* mutations failed to complement. Moreover, the phenotype displayed by *Pold1*^*tyrn/tm1b*^ embryos was more severe than that of the *Pold1*^*tyrn/tyrn*^ embryos (Fig. 2E), but milder than *Pold1*^*tm1b/tm1b*^ null mutants. These findings indicate that *Pold1*^*tyrn*^ is a hypomorphic allele and that perturbation of *Pold1* function is responsible for the phenotypes observed in *tyrn* mutants.

**Figure 2.**
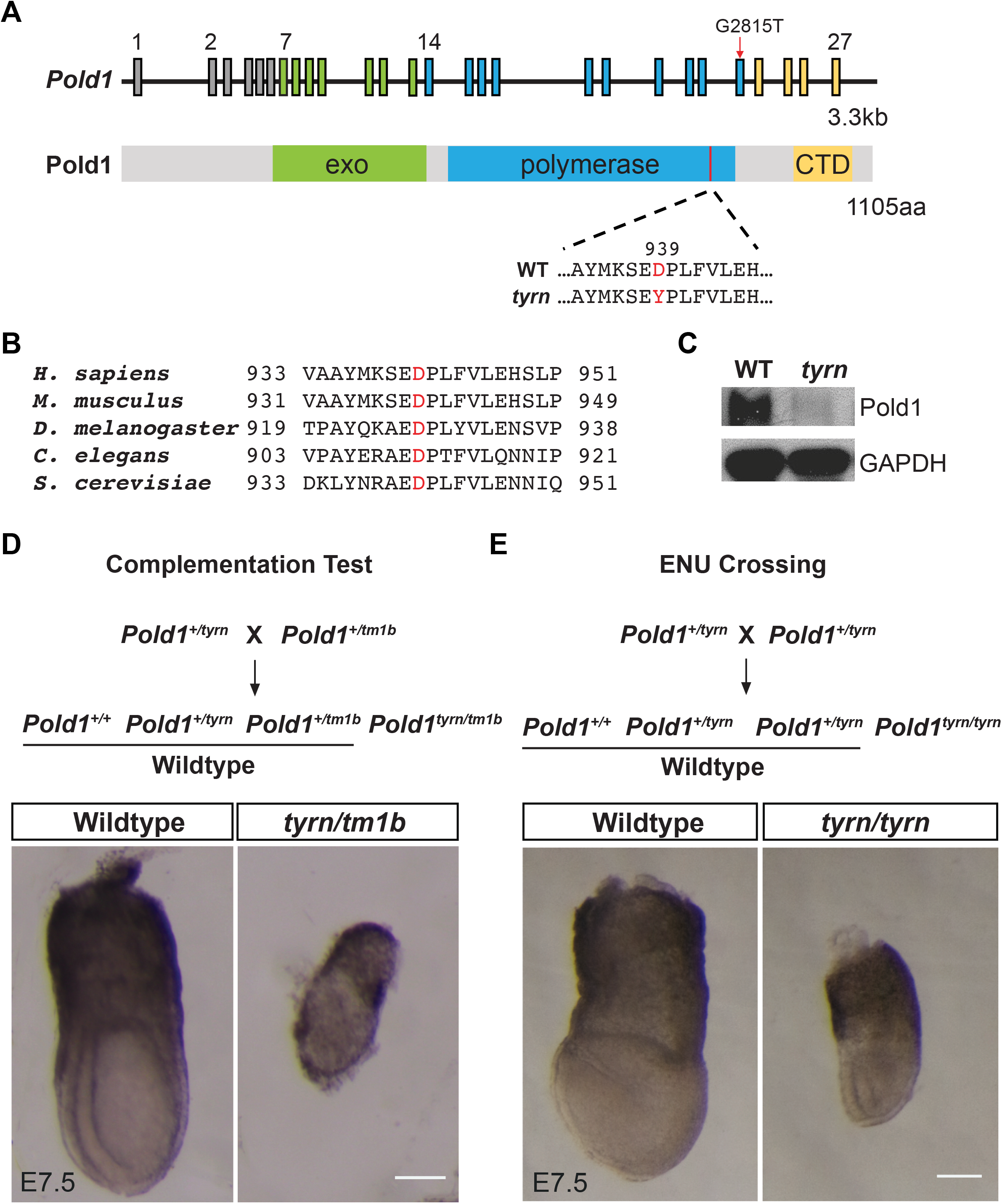
Identification of the *Pold1* missense mutation in *tyrn*. (A) Schematic diagrams of the murine *Pold1* locus in genome (upper panel) and of the Pold1 domain structure (lower panel). The G2815T nucleotide change (red arrow) was located in exon 23. The corresponding D939Y amino acid substitution was located in the DNA polymerase domain close to the C-terminal domain (CTD). (B) Multiple alignment of orthologous Pold1 amino acid sequences around D939Y among different eukaryotic organisms. The mutated aspartate residue was highly conserved. (C) Pold1 expression level in wildtype (left lane) and *tyrn* (right lane) embryos at E8.5 shown by western blot. (D) Complementation test crossing strategy (upper panel). Wildtype and *tyrn/tm1b* embryos acquired at E7.5 from the complementation test (lower panel). *tyrn/tm1b* embryos did not undergo gastrulation and were not able to survive past E7.5. (E) Crossing strategy to harvest homozygous mutants from heterozygous mice carrying *tyrn* allele (upper panel). E7.5 wildtype and mutant embryos (lower panel). n = 3 per genotype. Scale Bar = 100 μm.

### The *tyrn* mutation impairs DNA synthesis and cell proliferation

Based on the published cryo-electron microscopy structure of the human Pol δ (Lancey et al., 2020), the aspartate residue mutated in *tyrn* embryos resides in the thumb domain of the C-terminal DNA polymerase domain of POLD1 (Fig. 2A, Fig. S2A-C). Because of the important role of the thumb domain in stabilizing Pol δ at the primer-template junction during DNA synthesis (Jain et al., 2018), we hypothesized that the *tyrn* mutation may impair and reduce the DNA synthesis activity of Pol δ. We used a primer extension assay and mouse EdU labeling to test Pol δ polymerase activity *in vitro* and *in vivo*, respectively. For the primer extension assay, we modeled the Pold1 D939Y missense mutation in budding yeast Pol δ by introducing the equivalent mutation, D941Y, in the catalytic subunit, Pol3 (Devbhandari and Remus, 2020) (Fig. 3A). We purified wildtype Pol δ or Pol δ^D941Y^ (with Pol3^D941Y^) after overexpression in budding yeast (Fig. 3B). Analysis of the DNA synthesis products by denaturing gel-analysis reveals that both overall DNA synthesis and the level of full-length DNA products were significantly reduced in the presence of Pol3^D941Y^ when compared to the wildtype Pol δ (Fig. 3C-E). These data demonstrate that the D941Y substitution in Pol3, and by extension the corresponding D939Y substitution in Pold1, impairs the DNA polymerase activity of Pol δ, possibly by decreasing its processivity.

**Figure 3.**
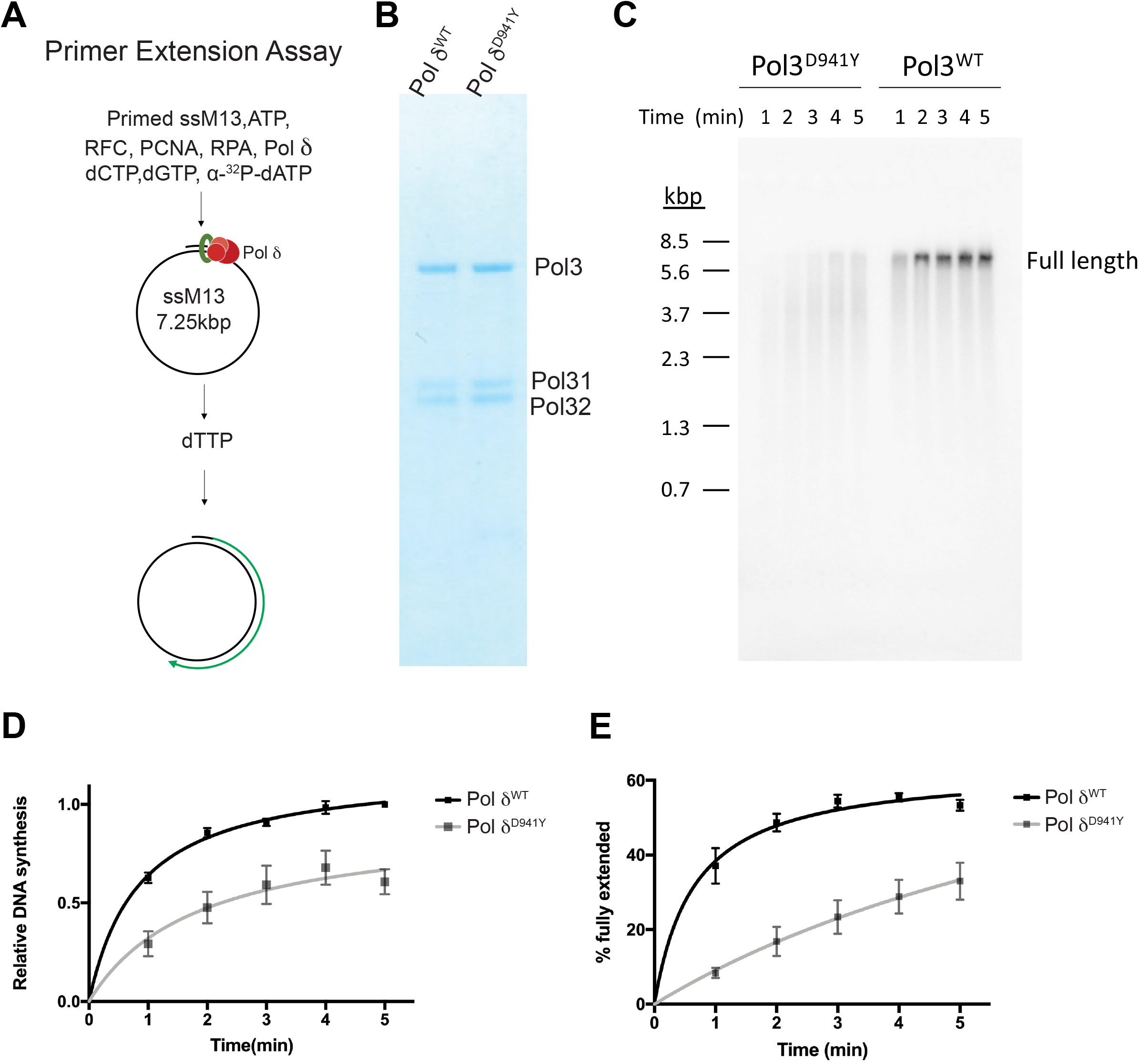
The effects of the *D941Y* mutation in *Pol3* on DNA synthesis. (A) Workflow of the *in vitro* primer extension assay. (B) Denatured protein gel showing the overexpressed yeast Pol δ^WT^ and Pol δ^D941Y^. The Pol3^WT^ and Pol3^D941Y^ corresponds to mouse Pold1^WT^ and Pold1^D939Y^, respectively. The Pol31 and Pol32 are the associated subunits in Pol δ corresponding to Pold2 and Pold3 in mouse. (C) *In vitro* primer extension assay showing reduction of DNA synthesis efficiency of Pol3^D941Y^, the yeast orthologue which harbors the corresponding D939Y mutation identified in mouse. (D) Nonlinear-fitted curves of total amount of new synthesized DNAs at different time points. The mean is represented by black squares (Pol δ^WT^) and grey squares (Pol δ^D941Y^). Error bars represent s.e.m.. For each time point, n=3 per genotype. (E) Nonlinear-fitted curves of the percentage of full-length circular DNAs among total products. The mean is represented by black squares (Pol δ^WT^) and grey squares (Pol δ^D941Y^). Error bars represent s.e.m.. For each time point, n=3 per genotype. Multiple Student’s *t* test, two-tailed, p<0.05.

To test if DNA synthesis is affected in *tyrn* embryos, we performed *in vivo* EdU labeling at E6.0, E6.5 and E7.0, the window of time during which the size difference between wildtype and mutant embryos emerges. We observed a significant reduction in EdU incorporation in mutants compared to wild types at all stages (Fig. 4A-B). Both wildtype and mutant embryos showed a substantial increase in cell number from E6.0 to E7.0, but *tyrn* embryos exhibited a slower growth rate starting from E6.5; by E7.0 mutant embryos had significantly lower numbers of cells compared to wildtype embryos (Fig. 4C). The reduction in cell number is not due to increased cell death; based on cleaved Caspase-3 staining, levels of cell apoptosis were similar between wildtype and mutant embryos during this period (Fig. S3). However, mutant embryos showed increased cell apoptosis at around E7.5, with apoptotic cells concentrated at the distal tip, the prospective location of the abnormal small head at E8.5 (Fig. 4D). Taken together, we conclude that the D939Y missense mutation impairs Pold1 polymerase activity, which, together with the reduced Pold1 protein expression in *tyrn* embryos, impedes cell proliferation and leads to a reduction in embryo size during gastrulation.

**Figure 4.**
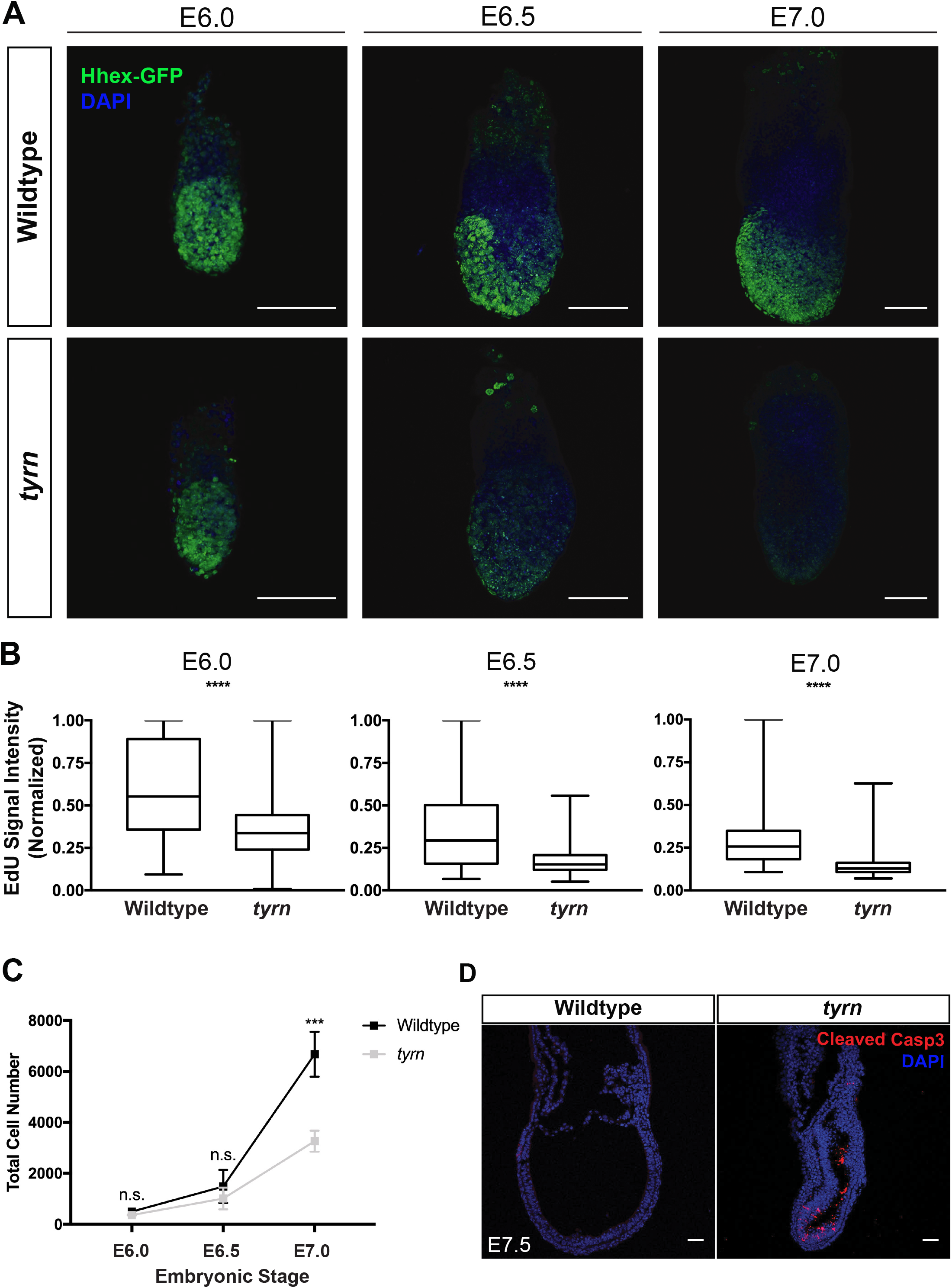
Analysis of DNA synthesis and cell proliferation in *tyrn* mutants. (A) EdU incorporation levels in wildtype and mutant embryos at E6.0, E6.5 and E7.0. *n*=3 embryos per genotype. Scale Bar = 100 μm. (B). Quantification of EdU signal intensity showing by box whisker plot. EdU signals of the embryos at the same stage was normalized to the maximum intensity in that stage. Total embryos quantified per genotype: *n*=3. Unpaired Student’s *t* test, two tailed, p<0.0001. Within box plot, the median is represented by the horizontal dividing line and the top and bottom of the box represents the seventy-fifth and twenty-fifth percentile, with the whiskers indicating the maximum and minimum points. (C) Curves of total cell number of wildtype and *tyrn* embryo at E6.0, E6.5 and E7.0 stages. n=3 embryos per genotype per stage. Unpaired Student’s *t* test, two-tailed, p<0.001. Within the curve, the mean is represented by the black squares (wildtype) and grey squares (*tyrn*). the error bars represent s.d.. (D) Immunofluorescence staining of Cleaved Caspase-3 in wildtype and mutant embryos at E7.5. n = 3 per genotype. Scale Bar = 50 μm.

### The *tyrn* mutation does not affect anterior visceral endoderm (AVE) positioning

The mispositioned head, and the abnormal morphology of E8.5 *tyrn* mutants, indicated that the A-P axis, although established, was shifted in orientation along the proximal-distal axis. Proper formation of the A-P axis depends on the anterior migration of a morphologically distinct population of VE cells from their position at the distal tip of the E5.5 embryo (Rivera-Perez et al., 2003); by E5.75-E6.0 this cell population will reach the anterior epiblast - extraembryonic ectoderm boundary and form the AVE (anterior visceral endoderm) (Srinivas et al., 2004). Multiple studies have shown that defects in AVE migration can lead to abnormal positioning of the A-P axis and mislocalized head phenotypes (Acampora et al., 2009; Clements et al., 2011; Kojima et al., 2014; Lu et al., 2001; Martinez-Barbera et al., 2000; Rossant and Tam, 2009). Therefore, we asked whether the AVE resided in its normal anterior position in *tyrn* mutants at E6.5. Although the *tyrn* mutants tended to be smaller at this stage, they were morphologically indistinguishable from wildtype littermates. WISH showed that *tyrn* mutants expressed two archetypal AVE markers, *Dkk1* (a Wnt antagonist) and *Cerl* (a Nodal antagonist), in a pattern comparable to that of wildtype embryos (Fig. 5A-B). We validated these findings by crossing in the Hhex-GFP transgene reporter to fluorescently label AVE cells in *tyrn* mutants (Rodriguez et al., 2001). Consistent with the *Dkk1* and *Cer1 in situ* hybridization results, we detected the Hhex-GFP expressing AVE cells at the anterior boundary between the embryonic and extraembryonic regions in *tyrn* embryos at E6.5 (Fig. 5C). These results indicate that AVE migration and A-P axis establishment proceed normally in pre-gastrulation stage *tyrn* mutants.

**Figure 5.**
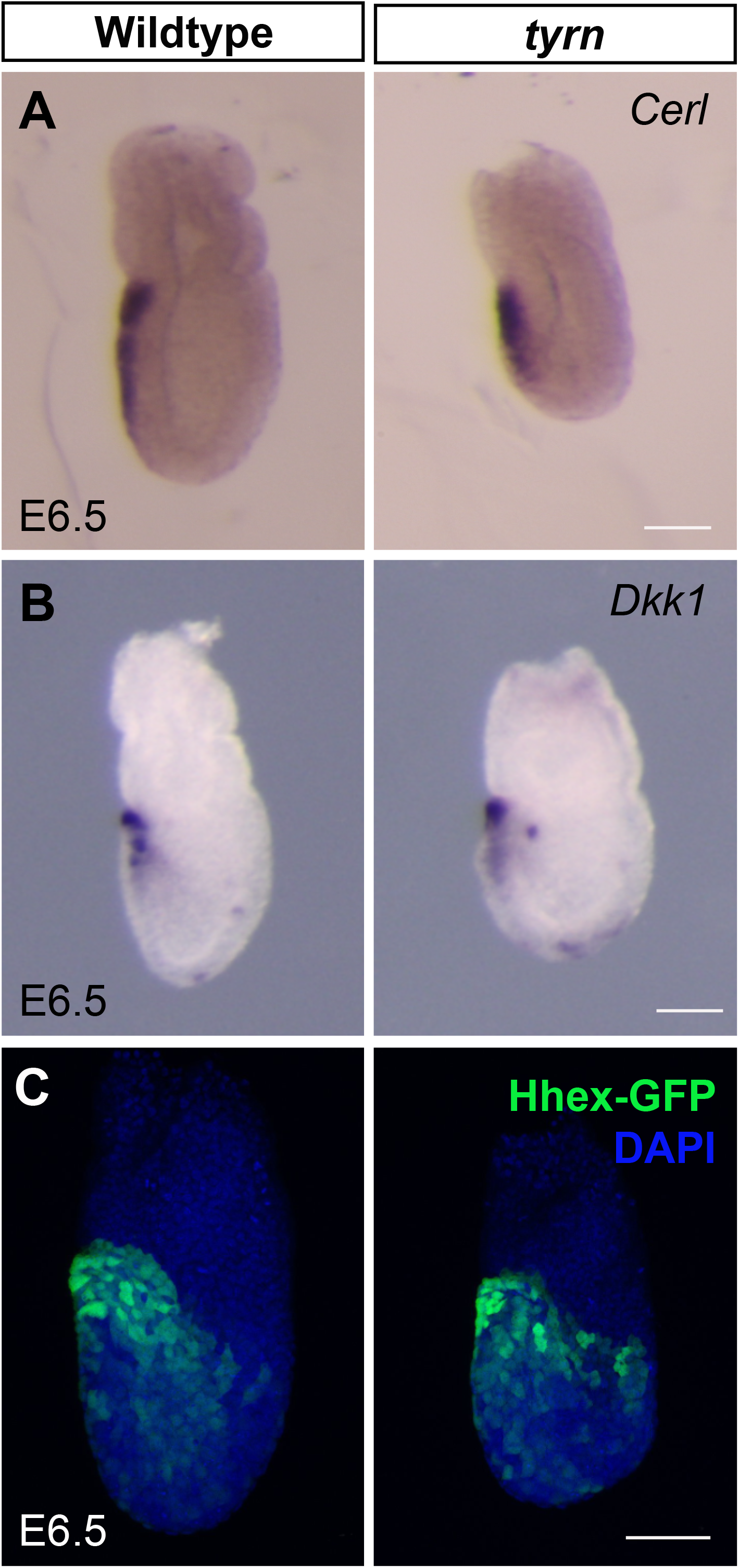
Position of Anterior Visceral Endoderm in *tyrn* mutants. *Cerl* (A) and *Dkk1* (B) were expressed in anterior visceral endoderm (AVE) in both wildtype and *tyrn* embryos at E6.5. (C) Hhex-GFP was expressed in AVE in both wildtype and *tyrn* embryos at E6.5. *n* = 3 embryos per genotype. Scale Bar = 50 μm.

### The *tyrn* mutation affects primitive streak extension and head position at E7.5

At E6.5, the AVE resided at the expected location in *tyrn* mutants; yet the position of the *tyrn* embryo’s anterior region was shifted toward the distal tip at E7.5. We examined the expression of multiple anterior-specific markers to visualize the organization of the anterior region in both wildtype and mutant embryos at E7.5. As depicted in Fig. 6A-B, wildtype embryos expressed Sox2 uniformly throughout the anterior epiblast, whereas the *tyrn* mutants expressed Sox2 in a discontinuous pattern, with more intense staining in the distal epiblast than in the proximal/anterior region. In the wildtype E7.5 embryo in the left panel of Fig. 6C & D, the Hhex-GFP transgene labels anterior definitive endoderm (ADE) as well as remaining AVE cells. In contrast, the Hhex-GFP staining resides predominantly in distal region of the E7.5 *tyrn* mutant, with no obvious A-P asymmetry (Fig. 6C-D, right panels). Similarly, cells expressing *Otx2*, a head organizer marker, lie distally in mutant embryos compared to their anterior position in wild types (Fig. 6E).

**Figure 6.**
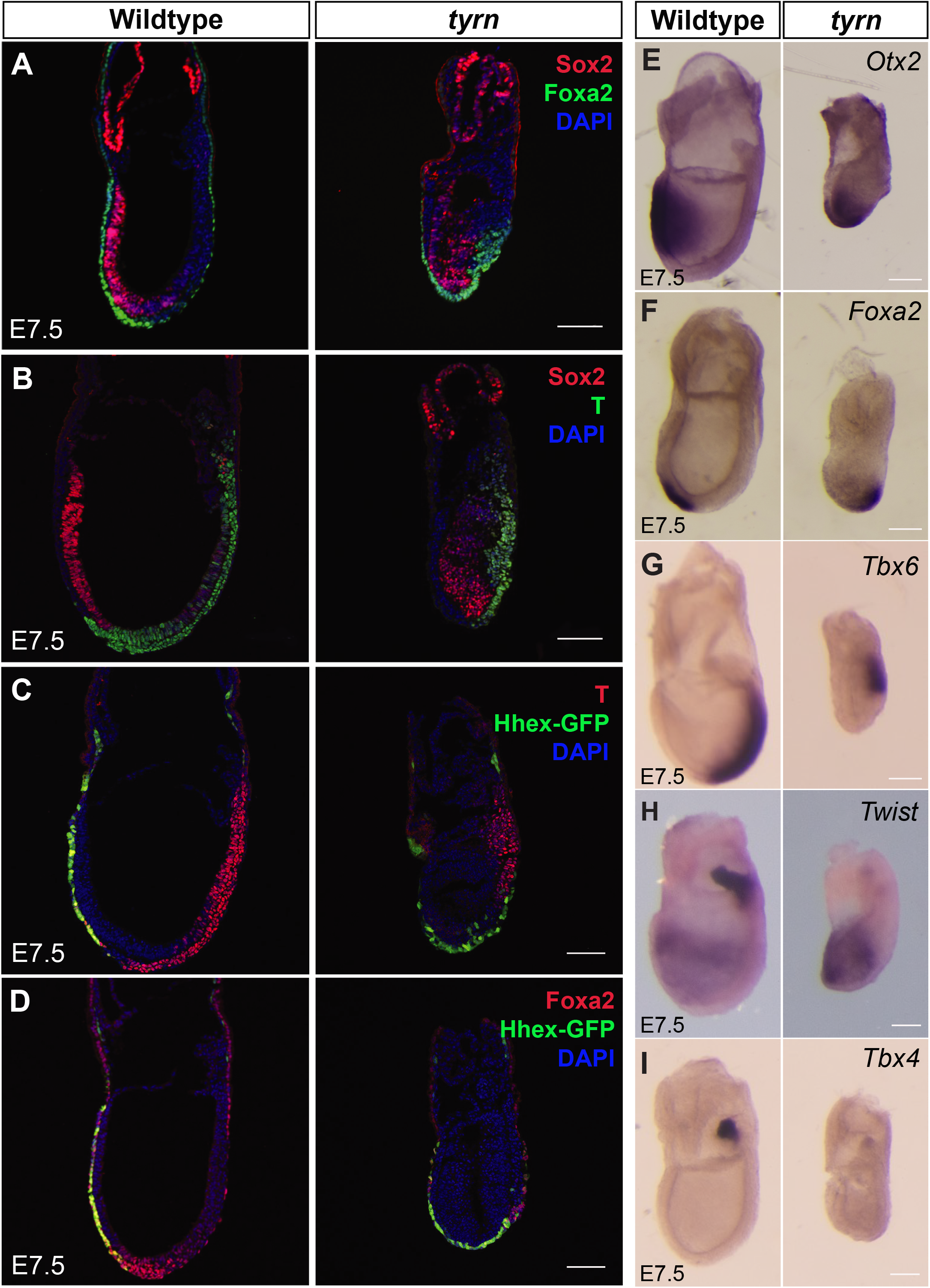
Characterization of primitive streak extension in *tyrn* mutants at E7.5. Co-staining of Sox2 with Foxa2 (A) or with T (B). Co-staining of with T (C) or Foxa2 (D). The anterior marker Sox2 was strongly expressed at the distal tip in mutants. Sox2 was also expressed in the ectoderm of chorion in both wildtype and mutant embryos. A short primitive streak was formed. AME and ADE cells emerged from the midpoint of posterior side. Sox2 labels the extraembryonic ectoderm of the chorion. (E) *in situ* hybridization staining of *Otx2* showed that the *Otx2* expression was restricted distally in *tyrn* rather than anteriorly seen in wild types. (F) *Foxa2* labels axial mesoderm and definitive endoderm cells. (G) Paraxial mesoderm marker *Tbx6* expression was restricted to the posterior, consistent with the T expression pattern. (H) *Twist* was first expressed in extraembryonic mesoderm and then strongly expressed in allantois. It was also expressed in embryonic mesoderm which leaves primitive streak. (I) *Tbx4* expression in allantois was strong in wild types but not seen mutant embryos at E7.5. n = 3 embryos per genotype. Scale Bar=100 μm.

The AVE serves as a transient anterior signaling center at ∼E6.5. As gastrulation progresses, AVE cells disperse into the extraembryonic/embryonic boundary and the anterior portion of the extraembryonic yolk sac (Lawson and Pedersen, 1987; Rivera-Perez et al., 2003; Shimono and Behringer, 2003; Tam et al., 2007). Meanwhile, the ADE and axial mesoderm (AME), both of which strongly express Nodal and Wnt antagonists (Arnold and Robertson, 2009), gradually migrate anteriorly to replace the AVE and become new sources of anterior signaling. We hypothesized that the abnormal orientation of the A-P axis might reflect distal positioning of AME and ADE. Fig. 6A, D, F examines the expression of *Foxa2*, a marker of AME and ADE, in wildtype and *tyrn* embryos at E7.5. Emerging *Foxa2*-labeled ADE and AME cells reside in the distal anterior region in wild types, but these cells sit more posteriorly in mutants.

The aberrant posterior positioning of ADE and AME, both derivatives of anterior primitive streak (APS) (Arnold and Robertson, 2009; Lawson, 1999; Lawson et al., 1991), suggested that primitive streak extension was defective in *tyrn* mutants. To compare the organization of the primitive between wildtype and mutant embryos, we performed immunofluorescence for T (Brachyury), a marker of nascent mesoderm emanating from the primitive streak, including AME (Fig. 6B-C). In E7.5 wildtype embryos, T staining showed that the primitive streak had extended to the distal tip and generated AME derivatives of the anterior primitive streak. In contrast, T staining of E7.5 *tyrn* embryos indicated that the primitive streak had extended only to the midpoint of the posterior side (Fig. 6B-C). Also pointing to impaired primitive streak elongation in *tyrn* embryos, WISH detected irregularities in the production and positioning of paraxial and extraembryonic mesoderm. Whereas expression of *Tbx6*, a marker of paraxial mesoderm, was found along the posterior side of the wildtype embryo, down to the distal tip, it was restricted to the posterior-proximal region in *tyrn*, consistent with a short primitive streak (Fig. 6G). *Twist* expression labels two distinct mesodermal populations at E7.5; anterior mesoderm precursors to cranial mesenchyme and extraembryonic mesoderm of the allantois (Bildsoe et al., 2009). Of note, the E7.5 *tyrn* mutant generated *Twist* expressing anterior mesoderm but lacked *Twist* expressing cells of the allantois (Fig. 6H). The mutant embryos also expressed greatly reduced levels of *Tbx4*, another marker of allantoic mesoderm (Fig. 6I). These findings indicate that *tyrn* mutants have very few extraembryonic mesoderm cells inside the allantois. Taken together, these results suggest that the impaired extension of the primitive streak in *tyrn* embryos causes defects in distribution of multiple mesoderm derivatives, including the distal positioning of the AME and ADE at E7.5. Mutant embryos continued with head morphogenesis and eventually formed a head at the distal tip with abnormal morphology.

## Discussion

The regulation of cell number plays an integral role in mouse embryo size determination during the pre-gastrulation stages. Earlier studies used embryo manipulation or pharmacological methods to alter cell numbers in pre- and early post-implantation mouse embryos. The findings showed, that prior to gastrulation, mouse embryos have the ability to sense and respond to an acute, global alteration in cell numbers and achieve normal size. Thus, robust size regulatory mechanisms must operate intrinsically in the mouse embryo to ensure proper size before the onset of gastrulation (Lewis and Rossant, 1982; Power and Tam, 1993). These classic studies raise many questions about when, in what cell populations, and how size regulatory mechanism function; questions that are not amenable to approaches involving the global loss or gain of cells especially for investigations around and after gastrulation. Fine-tuned genetic methods could allow manipulation of candidate components of the size regulatory process for extended period. Here, we assessed the impact on embryo growth and morphogenesis of a hypomorphic mutation that reduces levels of DNA proliferation at gastrulation stages. The *tyrn* mutants from our ENU mutagenesis screen provided an excellent time window for us to investigate the roles of embryo growth in gastrulation. The *tyrn* mutation disrupted the polymerase function of Pold1 and caused reduced cell proliferation *in vivo*. Mutant embryos did not merely exhibit a wildtype-like morphology with a proportionally reduced size, or a random shape with no underlying logic. Instead, it showed a siren-like morphology, with the head located closer to the distal tip rather than residing at the normal anterior region seen in the wild types. Further phenotypic analyses from E6.5-E7.5 stages revealed that the mispositioned head as well as the general abnormal morphology seen in *tyrn* embryos likely resulted from the defective primitive streak extension during gastrulation. The short primitive streak led to the distal positioning of the AME and ADE, which caused the orientation of the anterior-posterior axis to be shifted towards the distal-proximal axis (Fig. 7). The *tyrn* mutants showed remarkable impact of reduced cell proliferation during gastrulation. The impaired DNA synthesis machinery caused prolonged defects in cell proliferation, leading to a discoordination of embryo growth with lineage specification and tissue morphogenesis. Such discoordination resulted in embryos with abnormal morphology and incorrect orientation inside the decidua. These results indicate that the embryo growth needs to be highly coordinated with lineage specification and tissue morphogenesis for normal gastrulation.

**Figure 7.**
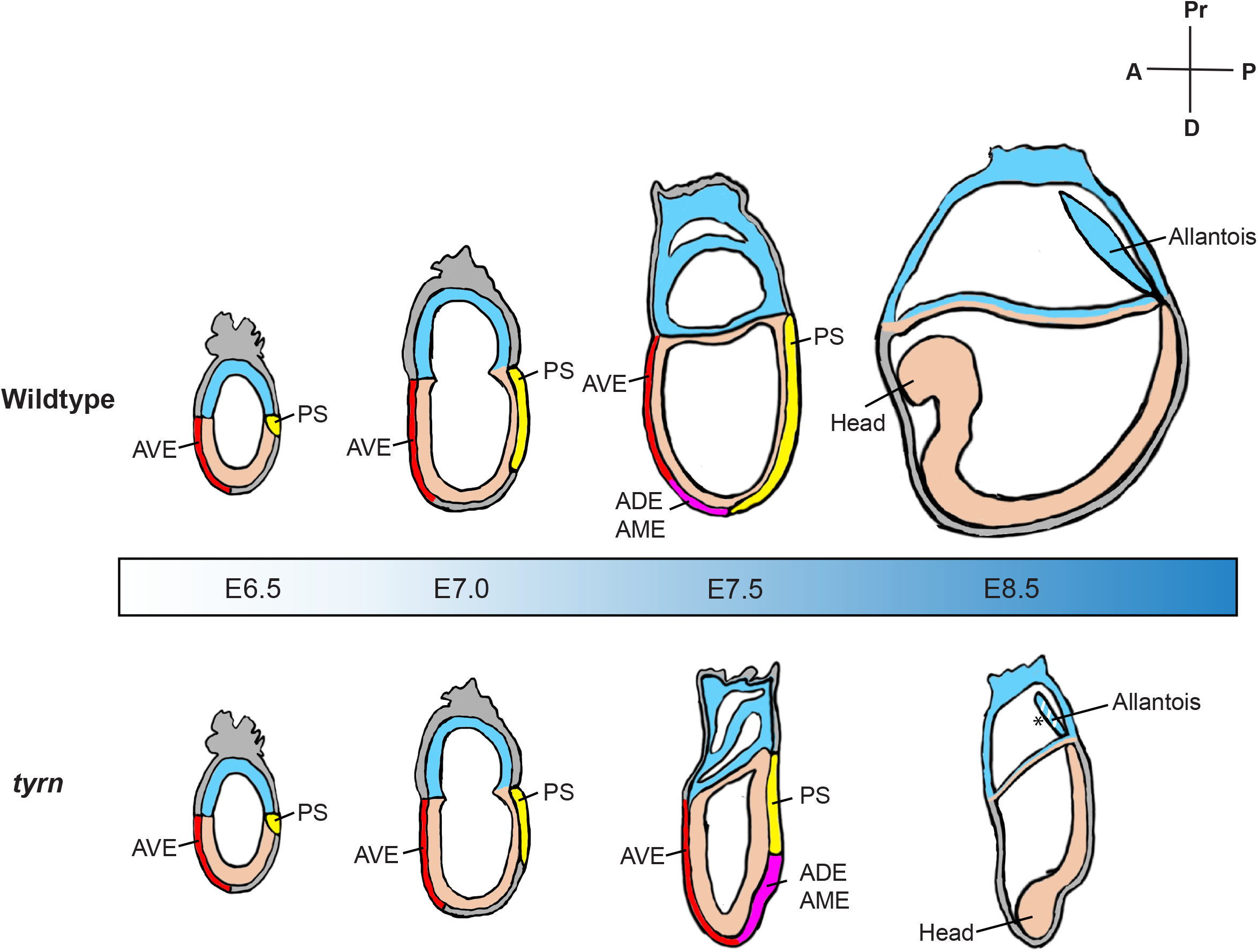
Developmental progression of wildtype and *tyrn* embryos between E6.5 and E8.5. At E6.5, wildtype and *tyrn* embryos are morphologically indistinguishable. AVE (red) is properly localized at the normal anterior region in both wildtype and mutant embryos. Gastrulation is initiated and primitive streak is formed (yellow). At E7.0. primitive streak elongation (yellow) in *tyrn* shows slight delay compared to wildtype embryos. Embryo size also starts to show difference although still not visually obvious. At E7.5, the primitive streak extension in *tyrn* was delayed. The AME and ADE emerged from the midpoint of the posterior side and did not migrate across the bottom point of the embryos (magenta). The distal tip in mutants becomes the anterior signal center and embryos consider the distal tip as the place for head induction and morphogenesis. At E8.5, the wildtype embryos are properly developed along normal A-P axis, whereas the mutant embryos exhibit a small, siren-like morphology with head pointing to the bottom of the embryo. The asterisk indicates the reduction of extraembryonic mesoderm cells in the allantois.

Our study suggests that mesoderm lineage allocation during gastrulation is affected by the global reduction of cell proliferation. Primitive streak is initially induced at the posterior-proximal pole where high levels of BMP, FGF, Wnt and Nodal signals present (Brennan et al., 2001; Ciruna and Rossant, 2001; Conlon et al., 1994; Huelsken et al., 2000; Mishina et al., 1995). The primitive streak elongates and extends to the distal tip as gastrulation progresses. The cell fate of mesoderm derivatives is determined by timing and positions of their ingression through the streak (Vincent et al., 2003; Winnier et al., 1995). The normal cell proliferation rate guarantees coordination of embryo size expansion and the signal gradient, enabling cells along the streak being exposed to different signal combination and contribute to different mesoderm subtypes. Such coordination was disrupted due to reduced cell proliferation: the primitive streak failed to fully extend, which could lead to aberrant signal gradients along the streak. We found that axial mesoderm, paraxial mesoderm subtypes were generated, albeit located more posteriorly. The greatly reduced allantoic mesoderm may be a consequence of altered posterior signal environment. It could also reflect a general developmental delay: the cell differentiation along the streak follows the anterior-posterior order and the allantoic mesoderm arises relatively late during gastrulation. Another possibility is that these extraembryonic mesodermal cells may originate from a highly proliferative cell population (Snow and Tam, 1979) that could be more sensitive to inhibition of DNA synthesis. These questions may be addressed in future through the perturbation of embryonic cell numbers in specific cell populations during gastrulation.

Cell number regulation occurs throughout different stages of embryonic development. The effects of cell number perturbation on tissue/organ size and morphogenesis can vary greatly based on developmental stages and tissue/organ types. The mandibular hypoplasia, deafness, progeroid and lipodystrophy (MDPL) syndrome, a multisystem disorder, has been associated with heterozygous in-frame deletion of Ser605 that causes loss of POLD1 activity (Elouej et al., 2017; Fiorillo et al., 2018; Sasaki et al., 2018; Weedon et al., 2013). The reported patients presented growth retardation, sensorineural deafness, loss of subcutaneous adipose tissue and insulin resistance, indicating that decreased POLD1 activity exerts organ specific effects. Other studies have also identified organ-specific responses to loss of tissue-specific progenitors. Removing the embryonic limb field in amphibians or chickens has no effect on the final limb size as the limb is capable of robust compensatory proliferation (Holder, 1981; Summerbell, 1981), and similar compensatory growth has been observed for the developing liver progenitors (Bort et al., 2006). In contrast, the final pancreas size is determined by the initial pancreatic progenitor pool and there is a lack of significant compensatory growth (Stanger et al., 2007). These studies demonstrate the importance of studying cell number regulation in diverse tissue and organ types at different stages of embryonic development. Our findings underscore the power of unbiased phenotype-based forward genetic screening approaches and highlight the value of using ENU-induced hypomorphic alleles to uncover new biological functions of well characterized genes such as *Pold1*. For decades, embryo size regulation has become the “forgotten classics” in the developmental biology field (https://thenode.biologists.com/forgotten-classics-regulating-size-mouse-embryo/research/). In future studies, one may use carefully chosen genetic models coupled with modern developmental biology techniques such as live imaging and single cell genomics to revisit this important topic in diverse developmental contexts beyond the early stages of embryogenesis.

## Materials and Methods

### Mouse Strains (*Mus musculus*)

The parental JM8.N4 mouse ES Cell Strain (strain origin: C57BL/6N, male, black coat, non-agouti, MGI ID: 4431772) (Skarnes et al., 2011) carrying one *Pold1*^*tm1a(EUCOMM)Wtsi*^ (*Pold1*^*tm1a*^) allele was imported from European Mouse Mutant Repository (EUMMCR) unit. The selected Pold1-G09 ES cell clone passed the karyotype check performed by Molecular Cytogenetics Core Facility, Memorial Sloan Kettering Cancer Center (MSKCC) and was free of mouse pathogens tested by MSKCC Center of Comparative Medicine & Pathology. Pold1-G09 ES cell clone was injected into the female B6(Cg)-Tyr^c-2J^/J (albino C57BL/6J, or B6-albino, non-agouti, The Jackson Laboratory) donor mice by MSKCC Mouse Genetics Core Facility. 14 out of the 19 pups alive were chimera. Male chimeras were crossed with albino FVB/NJ (FVB, containing homozygous dominant *agouti* locus *A*/*A*) females to generate heterozygous offspring carrying the *Pold1*^*tm1a*^ allele. Germline transmission was determined by the presence of agouti pups and genotyping. The *Pold1*^*tm1a*^ allele contains a *lacZ* cassette for genotyping. Mice carrying *Pold1*^*tm1b*^ (null) allele were generated by crossing *Pold1*^*tm1a/+*^ mice with *CAG-Cre* transgenic mice (The Jackson Laboratory) to remove the critical exons between exon 3 to exon 10. The *lacZ* cassette remained in *Pold1*^*tm1b*^ allele and was used for genotyping. The Hhex-GFP strain was a gift from Anna Katarina Hadjatonakis (Rodriguez et al., 2001). Mice that were 8-16 weeks old were used to generate E6.5 to E8.5 embryos. Analysis of the mutant phenotype was performed in the FVB genetic background or FVB-B6 mixed background. Mice were housed and bred under standard conditions in accordance with Institutional Animal Care and Use Committee (IACUC) guidelines. The MSKCC IACUC approved the experiments.

### ENU Allele Isolation, Sequencing and Genotyping

The *tyrn* allele was generated by mouse ENU mutagenesis screens using C57BL/6J males and identified based on its embryonic phenotype at E8.5, as previously described (Garcia-Garcia et al., 2005). To obtain genomic DNA, E8.5 mutant embryos were pooled into 3 sample groups and snap frozen on dry ice. Gentra Puregene kit (QIAGEN) was used for genomic DNA extraction of mutant embryos. Whole-exome sequencing was performed at the MSKCC Integrated Genomics Operation. Exome capture was performed using SureSelectXT kit (Agilent Technologies) and SureSelect Mouse All Exon baits (Agilent Technologies). An average of 100 million 75-bp paired reads were generated. Sequencing data analysis was performed using the methods described in (Jain et al., 2017). To obtain a list of potential phenotype-causing lesions, variants were filtered further to only include those that 1) were not found in dbSNP, 2) were not found in samples from other lines sent for this sequencing batch, 3) were shared among all 3 mutant samples, 4) were with coverage (sequencing read depth) of at least 6, 5) were homozygous, 6) were exonic only. 11 variants were found in 8 genes/annotations, 6 nonsynonymous, of which *Pold1* was one: *Pold1*: NM_011131: exon23: c. G2815T: p.D939Y. After examining the known phenotypes of published alleles or functional annotations of these candidate genes, we re-examined non-exonic and heterozygous mutations and eliminate read depth threshold to explore other possibilities. After going through the whole dataset (developed by Devanshi Jain from Scott Keeney lab)(Jain et al., 2017) containing all our exome sequencing submissions over the years to catalog universal variants we could eliminate, plus additional analysis with the above criteria, we confirmed that the sole candidate variant was in *Pold1*. The *tyrn* allele has a single G to T transversion in the 2815 nucleotide position of exon 23 in *Pold1* coding sequence (G2815T), causing a missense mutation (in the amino acid position 939 aspartic acid to tyrosine, D939Y). The G2815T mutation created a RsaI restriction site used for genotyping.

### Complementation Test

*Pold1*^*+/tm1b*^ females were crossed with *tyrn/+* males to produce embryos. Embryos at E7.5 and E8.5 stages were retrieved and grouped into wild types and mutants based on phenotypes. No mutant embryos were found at E8.5 stage. The genotypes of E7.5 embryos matched their phenotypes: Embryos with mutant phenotype were *Pold1*^*tyrn/tm1b*^; *Pold1*^*+/+*^, *Pold1*^*+/tyrn*^ *and Pold1*^*+/tm1b*^ showed wildtype phenotypes.

### Embryo Harvesting and Dissections

Pregnant FVB female mice bearing embryos at E6.5-10.5 stages were euthanized by cervical dislocation. Uterus were taken out for dissection based on approved mouse protocol. Embryos were dissected from decidua inside uterus in cold 0.4% Bovine Serum Albumin (Sigma-Aldrich) in Phosphate-buffered Saline (PBS) using fine forceps and Leica dissection scope. Embryos were fixed in 4% paraformaldehyde (PFA) on ice for at least 4 hours and washed with PBS for 5 minutes X 3 times at room temperature (RT). Fixed embryos were stored in PBS at 4°C for future experiments.

### *In Situ* Hybridization

Fixed embryos were sequentially dehydrated in 25, 50%,75% methanol-depc PBS and 100% methanol and stored in -20°C. Embryos were taken out from methanol and sequentially rehydrated in 75%, 50%, 25% methanol-depc PBS before experiments. *In situ* Hybridization were performed following standard protocol (Eggenschwiler and Anderson, 2000). Briefly, after rehydration, embryos were incubated in 1 μg/mL Proteinase K/PBS solution for 3 to 7 min and hybridized in hybridization solution with RNA probes at 70°C overnight. A series of washes in 2X SSC and MAB solutions were performed the next day. The embryos were incubated in 1:10000 anti-Digoxigenin antibody (Roche) in blocking buffer (1% blocking reagent [Roche], 10% heat-inactivated goat serum, 0.1% Tween-20) in PBS overnight and washed extensively in PBS+0.1% Tween-20. Embryos were incubated in BM purple solution (Roche) at RT protected from light until the purple color was developed.

### EdU labeling

EdU (5-ethynyl-2’-deoxyuridine) powder (Invitrogen) was dissolved in sterile PBS into a working concentration of 2.5 mg/ml. Mice were weighed and injected with EdU solution (25mg/kg) intraperitoneally. Embryos were harvested 2 hours after injection and fixed in 4% PFA overnight. Fixed embryos were washed with PBS+3% BSA twice and then permeabilized in PBS+0.5%Triton X-100 at RT for 20 minutes. Embryos were washed in PBS+3% BSA after permeabilization. Embryos were incubated with reaction cocktail made from Click-iT™ EdU Cell Proliferation Kit for Imaging, Alexa Fluor 637 dye kit (Invitrogen) at RT for 30 minutes, protected from light. Cocktail was removed after incubation and embryos were washed in PBS+3% BSA twice.

### Immunofluorescence and Confocal Microscopy

Fixed embryos were store in PBS at 4°C before use. Embryos for cryosection were dehydrated in 30% Sucrose-PBS at 4°C overnight. Embryos were embedded in Tissue-Tek^®^ O.C.T Compound (Sakura Finetek) and frozen on smashed dry ice immediately after embedding. Embedded embryos were stored in -80°C before use. Embryos were sectioned in 10 μm using Leica CM1520 Cryostat. Section slides were stored in -80°C before use. For whole-mount staining embryos, embryos were permeabilized with PBS+0.5%Triton X-100 for 1 hour at RT. For section staining, slides were dried for 30 minutes and incubated in blocking buffer (0.1% TritonX-100, 1% heat-inactivated donkey serum [Gemini: Bio-produces] in PBS) for 1 hour at RT. Primary antibodies were diluted with the optimized diluting ratio in blocking buffer: Foxa2 (Abcam Cat# ab108422, RRID:AB_11157157,1:300), T (Cell Signaling Technology Cat# 81694, RRID:AB_2799983,1:400), Cleaved Caspase-3 (Cell Signaling Technology Cat# 8202, RRID:AB_1658166,1:300), Sox2 (R and D Systems Cat# AF2018, RRID:AB_355110,1:300). Embryos and slides were incubated with primary antibodies at 4°C overnight on rotator. Embryos and slides were washed with PBS for 10 minutes X 3 times the next day. Section slides or embryos were incubated in blocking buffer containing specific secondary antibodies (Invitrogen,1:500) and DAPI (1:1000) for 2 hours at RT. For whole mount staining embryos, embryos were washed in PBS for 5 minutes X 3 times and incubated in FocusClear (CelExplorer.Co) for 20 minutes at RT, protected from light. Embryos were mounted with MountClear (CelExplorer.Co) and stored at 4°C. For cryosection staining, slides were washed in PBS for 5 minutes X 3 times and mounted with ProLong Gold Antifade Mountant (Thermo Fisher Scientific) and stored at 4°C. Both embryos and sections were imaged using Leica SP8 inverted laser scanning confocal microscope. Confocal images were reconstructed using Fiji (ImageJ) open-source image processing software.

### Immunoblotting

E8.5 wildtype and mutant embryos were harvested and pooled separately and stored in -80°C before experiments. Tissues were homogenized in cold lysis buffer (0.1% NP-40, 50mM Tris-HCl [pH 7.2], 250mM NaCl, 2mM EDTA, phosphatase inhibitor mixture I and II [Calbiochem] and one tablet of Minicomplete [Roche] per 10ml) on ice. Lysate was left on shaker at 4°C for another 30 minutes. Lysate was centrifuged at maximum speed (12000 rpm) to collect supernatant. Protein concentration was determined through BSA-based protein assay using Quick Start Bradford 1X Dye Reagent (BIORAD) and adjusted to 2ug/ul. Samples were mixed 1:1 with 2X SDS loading buffer and denatured at 95°C for 5 minutes. Samples were loaded with equal amount onto 8% SDS-PAGE gels and ran for 2 hours under 150V in 1X SDS at RT. Proteins were transferred to PVDF membranes under 15V overnight at 4°C. Membranes were incubated in blocking buffer containing (TBST+5% BSA) for 1hour at RT and incubated with primary antibody in blocking buffer at 4°C overnight: POLD1 (Abcam Cat# ab168827,1:500); GAPDH (Santa Cruz Biotechnology Cat# sc-32233, RRID:AB_627679, 1:1000). Membranes were washed with TBST and incubated with specific secondary antibodies for 1 hour at RT. Membranes were washed with TBST and incubated with Pierce ECL Western Blotting Substrate (Thermo Fisher Scientific) for 5 minutes and were film-exposed to show target bands.

### Primer Extension Assay

Primer extension reactions were performed at 30°C in polymerization buffer (25 mM Tris-HCl pH 7.5, 8 mM Magnesium Acetate, 5 mM Potassium Glutamate, 5% Glycerol). Purified proteins used in the primer extension assays were purified as previously described (Devbhandari and Remus, 2020). DNA template for the assay was generated by annealing a primer (5′-CCCAGTCACGACGTTGTAAAACG-3′) to M13mp18 single-stranded DNA (New England Biolabs, N4040S). The assay was initiated by incubation of 1nM of DNA template with 1 mM ATP, 1 mM DTT, 80 μM dATP, 80 μM dGTP, 80 μM dCTP and 400 nM of RPA for 5 minutes. PCNA and RFC were then added to 70 nM and 4 nM, respectively, and incubation was continued for 5 minutes. Then, 33 nM of α−^32^P-dATP (3,000 Ci per mmol) and 4 nM of either Pol δ^WT^ or Pol δ^D941Y^ was added to the reaction resulting in a primer extension by 9 base pairs (due to lack of dTTP). After 5 min, 80 μM dTTP were added to the mix for synchronous primer extension. Equal volume aliquots of this reaction (18 μl) were removed from the master reaction (100 μl) at indicated times and stopped by adding EDTA and SDS to final concentrations of 40 mM and 0.25%, respectively. Products were fractionated on a 0.8% alkaline agarose gel (30 mM NaOH and 2 mM EDTA), dried and imaged using Typhoon FLA 7000. Quantification of the gel images was performed using the ImageJ.

### Quantitation of Total Cells, EdU Signal Intensity, EdU-Positive Cells

E6.0, E6.5 and E7.0 whole mount EdU-labeled embryos were imaged using Leica SP8 inverted laser scanning confocal microscope and confocal z stacks of embryos were generated. Mutant and wildtype embryos of the same stage were imaged under the same conditions. For each stage, 3 embryos per genotype were imaged for quantification. Optical sections of each embryo were imported into Imaris (version 9.5, Oxford Instruments), and 3D-reconstitution along Z-axis was performed for data analysis. We used spots function in Imaris to automatically segment all DAPI-positive cells (both embryonic and extraembryonic tissues) to obtain a total cell count for each embryo. EdU signal intensity of each segmented cell was automatically measured and the background signal was subtracted. Cells with EdU signal above 10 arbitrary unit were considered EdU positive and counted as EdU positive cells by the software. We used Prism 9 (GraphPad) to perform normalization of EdU signal intensity. At each stage, the EdU signal of each EdU positive cell was normalized to the maximum EdU signal intensity. Data points were presented as box plot to show the distribution of EdU signal intensity among EdU positive cell population. Within box plot, the median is represented by the horizontal dividing line and the top and bottom of the box represents the seventy-fifth and twenty-fifth percentile, with the whiskers indicating the maximum and minimum points. Two-tailed Student’s *t* test was performed to evaluate the significance of all measurements.

### Statistics and Graphs

Embryo images n = 3 for all the experiments. For primer extension assay, 3 biological replicates were performed. We used Prism 9 to perform 2-tailed Student’s *t* test to evaluate the significance of all measurements and generated box plots and non-linear fitted curves. For the structure of human DNA polymerase δ, the original structure was imported and processed in PyMOL (The PyMOL Molecular Graphics System, Version 1.2r3pre. Schrödinger, LLC)

## Data availability

The structure of human DNA polymerase delta holoenzyme was pulled from protein data bank (PDB). PDB DOI: 10.2210/pdb6TNY/pdb. EM Map EMD-10539: EMDB EMDataResource.

## Primer information

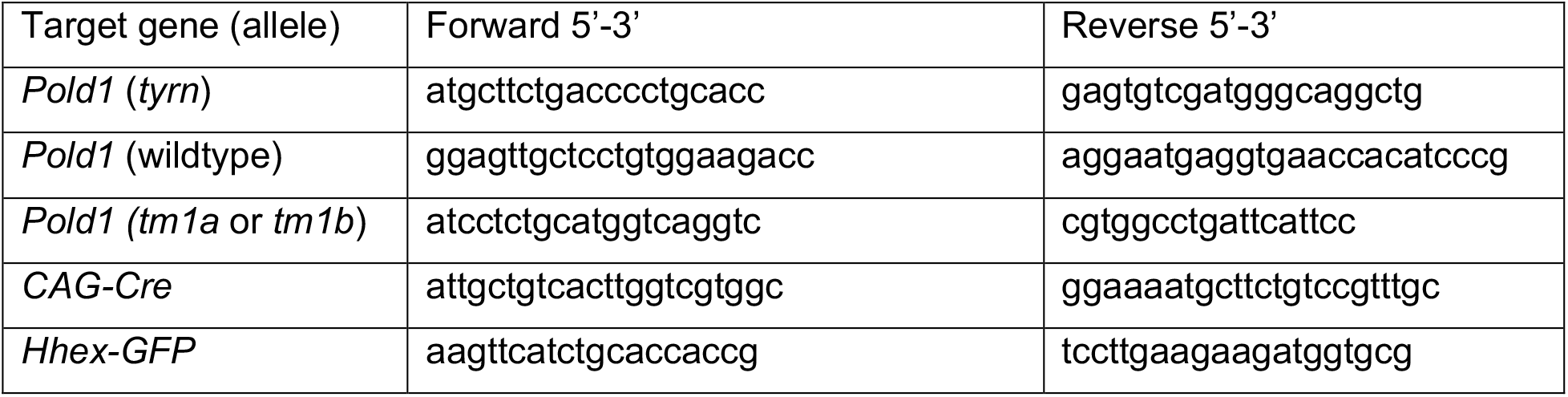

## Acknowledgments

The authors thank Edward Espinoza for *Pold1* cDNA preparation and validation of *Pold1* G2815T mutation, Dr. Devanshi Jain for sharing the methods of data analysis. The authors thank MSKCC Integrated Genome Operation for performing whole-exome sequencing and exome-wide SNP dataset, MSKCC Molecular Cytogenetics Core Facility for karyotype check, MSKCC Center of Comparative Medicine and Pathology for pathogen exam, MSKCC Mouse Genetics Core Facility for ESC clone injection, MSKCC Molecular Cytology Core Facility (Dr. Boyko, Mr. Tipping, Mr. Feng) at for assistance with confocal imaging and image processing. The authors thank Dr. Anna-Katerina Hadjantonakis for providing Hhex-GFP mouse strain, Jonathan Pai from Hadjantonakis’ lab for methods of clearing embryos. The authors thank Dr. Jennifer Zallen and Dr. Maria Jasin for helpful suggestions on this project.

## Author contributions

T.C. and K.V.A. conceived the study and designed the experiments. T.C., D.H. and E.L. interpreted the data and wrote the manuscript with inputs from other authors. K.V.A. and H.A. performed the ENU mutagenesis screen. T.C. maintained mouse lines, performed animal crossing, genotyping, dissection, *in situ* hybridization, immunofluorescence staining, confocal imaging, cloning, immunoblotting and data analysis. H.A. performed genomic DNA preparation and whole-exome sequencing data analysis. D.R. supervised and S.D. performed *in vitro* primer extension assays and quantifications.

## Figure Legends

**Supplementary Figure 1. Summary of *Pold1* mutant mice and crossing strategy of generating *Pold1* null allele**. (A) Schematic domain structure in mouse Pold1. The red bars represent mutations generated in previous genetic studies documented in the literature. The yellow bar denotes the missense mutation identified from ENU mutagenesis screen in this study. (B) Structure of the *Pold1* tm1a (gene trap) allele and the tm1b (null) allele generated after CAG-Cre-mediated recombination. Exons are represented in grey vertical blocks.

**Supplementary Figure 2. Cryo-EM structure of human POLD1 associated with DNA duplex and PCNA**. (A) Structure of the POLD1-DNA-PCNA complex colored by the domain and sequence of the DNA. (B) Zoom-in image showing the POLD1 thumb domain (red). The mutated Asp 941(D941) corresponsive to mouse Asp 939 (D939) residue was highlighted in lemon. (C) Zoom-in image showing the POLD1 thumb domain. The mutated Asp 941(D941) residue was shown as lemon stick. The protein structure information was extracted from protein data bank (PDB). Structure ID: 6TNY. PDB DOI: 10.2210/pdb6TNY/pdb. The original protein structure was adapted using PyMOL to deliver information relevant to this study.

**Supplementary Figure 3. Cell apoptosis in E6.5 embryos**. Whole mount staining of E6.5 wildtype and *tyrn* embryos. Cleaved Caspase-3 (Red) stained for apoptotic cells. DAPI (blue) stained nucleus of all cells. n = 3 per genotype. Scale Bar = 100 μm.

